# STARTER : A Stand-Alone Reconfigurable and Translational OoC Platform based on Modularity and Open Design Principles

**DOI:** 10.1101/2025.05.13.653800

**Authors:** A. Paul, E. R. Safai, L. E. de Heus, A.R. Vollertsen, K. Weijgertse, B. de Wagenaar, H. E. Amirabadi, E. van de Steeg, M. Odijk, A.D. van der Meer, J. Loessberg-Zahl

## Abstract

Organ-on-Chips (OoC) have the potential to revolutionize drug testing. However, the fragmented ecosystem of available OoC systems leads to wasted resources and collaboration barriers, slowing uptake. To address this, there is a need for OoC platforms based on interoperability standards, modularity, and reconfigurability. Technology platforms based on open designs would enable seamless integration of diverse OoC models and components, facilitating translation. Our study introduces a modular microfluidic platform that integrates swappable modules for pumping, sensing, and OoCs, all within the ANSI/SLAS microplate footprint. Sub-components operate as microfluidic building blocks (MFBBs) and can interface with the demonstrated Fluidic Circuit Board (FCB) universally as long as the designs adhere to ISO standards. The platform architecture allows tube-less inter-module interactions via arbitrary and reconfigurable fluidic circuits. We demonstrate two possible fluidic configurations which include in-line sensors and furthermore demonstrate biological functionality by running both in-vitro and ex-vivo OoC models for multiple days. This platform is designed to support automated multi-organ experiments, independent of OoC type or material. All designs shown are made open source to encourage broader compatibility and collaboration.

## 1 Introduction

Organ-on-Chips (OoCs) are advanced cell culture systems within controlled microfluidic environments. OoC research aims to replicate human physiological conditions more accurately for disease modelling and drug testing. Since their inception in 2010, OoC systems have evolved to include multi-organ systems and enhanced physiological realism, needing further capabilities in environmental control, sensing, actuation, and automation.^[1,2]^ With the increasing complexity of OoC systems, there is a strong emerging need for flexible, general architectures that can accommodate a variety of biological models and experimental setups. While both academic and industrial efforts have addressed the need for integrated micro-physiological systems, the lack of coherence leads to trade-offs in model complexity, throughput, and interoperability.^[3]^ This incompatibility impedes the development of innovative OoC models and hampers their operation and implementation in relevant end-user settings.^[4]^ Therefore, platforms based on modular design and standardized integration principles, such as those seen in the electronics industry, will be necessary to promote rapid innovation and future impact of OoC models.

Typical OoC studies rely on peripherals for functionalities like perfusion, parallelization, and sensing. However, traditional OoC systems often use peripherals from closed commercial ecosystems which include application-specific functionalities^[5]^. For example, platforms have been designed to cater to a particular OoC model ^[6-8]^, incorporate multiple OoCs on a single chip through monolithic integration^[9-11]^, support transwell-based models by employing specialized interfacing manifolds on fluidic boards^[12,13]^ or even offer fully integrated and self-contained solutions^[14-17]^. The diverse integration strategies adopted by these companies adds complexity and cost to the process of designing cross-compatible modules. Moreover, these highly specific and gatekept solutions limit translation across labs unless all labs make a similar investment in the same ecosystems, thereby increasing the capital required to conduct OoC research. While these closed ecosystems can offer a higher level of user-friendliness, the lack of modularity and inter-compatibility creates its own barrier to entry, especially for researchers with unique experimental needs.

Several custom platforms have been shown to offer more flexibility in integrating varying OoC models and sensors. However, these platforms typically offer low-throughput and limited external compatibility. For example, platforms have explored plug-and-play connections^[18-20]^, but the diversity of commonly-used microfluidic interconnection methods typically limits the use of the platforms to the founding labs. Multiplexed systems improve throughput^[21,30]^ but still rely on expensive peripherals and complex control systems, limiting scalability and broad adoption across research labs. As a result, despite various novel and creative solutions^[22-24]^, a universal integrating platform is yet to be reported. Standardized design guidelines are a first step towards universal interoperability, generating the opportunity to unite users, developers, and manufacturers in shared use of a custom OoC platform that has extended reach in the field. The ISO 22916:2022 ‘Microfluidic Devices’ standard provides a coherent framework to enable interoperability between microfluidic components^[25]^ from different sources. This paves the way to address the need for universal modular multi-OoC platforms that can perform automated experiments within commonly used lab workflows.^[26]^

One example of implementation of ISO 22916 relies on the concept of utilising a Fluidic Circuit Board (FCB) with the ANSI/SLAS standardized footprint^[27]^ of a microplate as the central fluidic manifold. Individual Microfluidic building blocks (MFBBs) with ISO-compliant footprints and port positions are connected to the FCB, resulting in a modular integrated platform. Dekker et al. have shown an initial catalogue of commercial components converted into ISO-compliant MFBBs.^[28]^ In the application domain of OoC, this concept of integrating standardized MFBBs and FCBs resulted in an open platform termed the ‘Translational Organ-on-Chip Platform (TOP)’^[3]^. Even though the ISO standards were not always strictly followed in these earlier platforms, the first conceptual implementations of TOP have demonstrated strong promise. For example, Vivas et al. demonstrated how commercial components can be combined with a custom Heart-on-Chip device for automated perfusion through an FCB^[29]^. De Graaf et al. demonstrated perfusion of 12 vessel-on-chips (VoC) on a FCB with controlled wall shear stress and circumferential strain^[30]^. Vollertsen et al. used similar interconnection and design principles with multiple highly multiplexed chips to demonstrate 192 individually controlled cell culture experiments on a modular platform.^[31]^

In all of the initial implementations of TOP, external peripheral equipment was still required to control the flow and pneumatic actuators. Moreover, they relied on a ‘hard-wired’ single fluidic circuit, which limited flexibility in experimental design. To enable broader uptake of TOP, it will be important to provide a set-up that does not rely on external peripheral equipment, is compatible with existing tissue culture workflows and can be adapted for use in many different applications. In short, we identify a strong need for a standalone, reconfigurable platform that supports a broad range of OoCs, sensors, and actuators and explicitly invites stakeholders’ adoption in the field.

This work presents STARTER - a TOP-based, standardized, modular, and reconfigurable microfluidic platform that integrates OoCs, sensors, reservoirs, and pumps from diverse manufacturers and materials. We demonstrate that the platform is standalone, combining three OoC devices with pumping and in-line sensing, all within a microtiter plate footprint. The platform’s architecture includes a ‘routing block’ that allows for complete reconfiguration of the fluidic circuits, enabling parallel, combined, or custom perfusion of the OoC devices. By adhering to ISO 22916, the platform ensures that compliant MFBBs can interface universally, whether commercially available or fully custom-designed. The specific port pitches and footprints of the MFBBs used in this work are openly documented. All design files for STARTER are publicly available in an open-source library, allowing the community to reproduce or build upon this work. This open-source approach aims to foster collaboration, to align research efforts in OoC, and to encourage wider industry adoption of standardized approaches.

## 2 Design

The STARTER platform is designed to be a standalone and flexible implementation of TOP to facilitate realization of a more diverse set of applications. This is achieved by dictating the fluidic circuits using the architecture of the FCB and a specific MFBB we refer to as ‘routing block’. The ‘routing block’ serves as a central hub where channels from all other modules are connected as required (Figure 1b). This strategy offers freedom in deciding interactions between various MFBBs and reconfigurability of the whole system by replacing a single MFBB. Together, the FCB and the routing block lay the foundation of a flexible architecture that facilitates multiple operation modes even with a fixed set of MFBBs.

**Figure 1.**
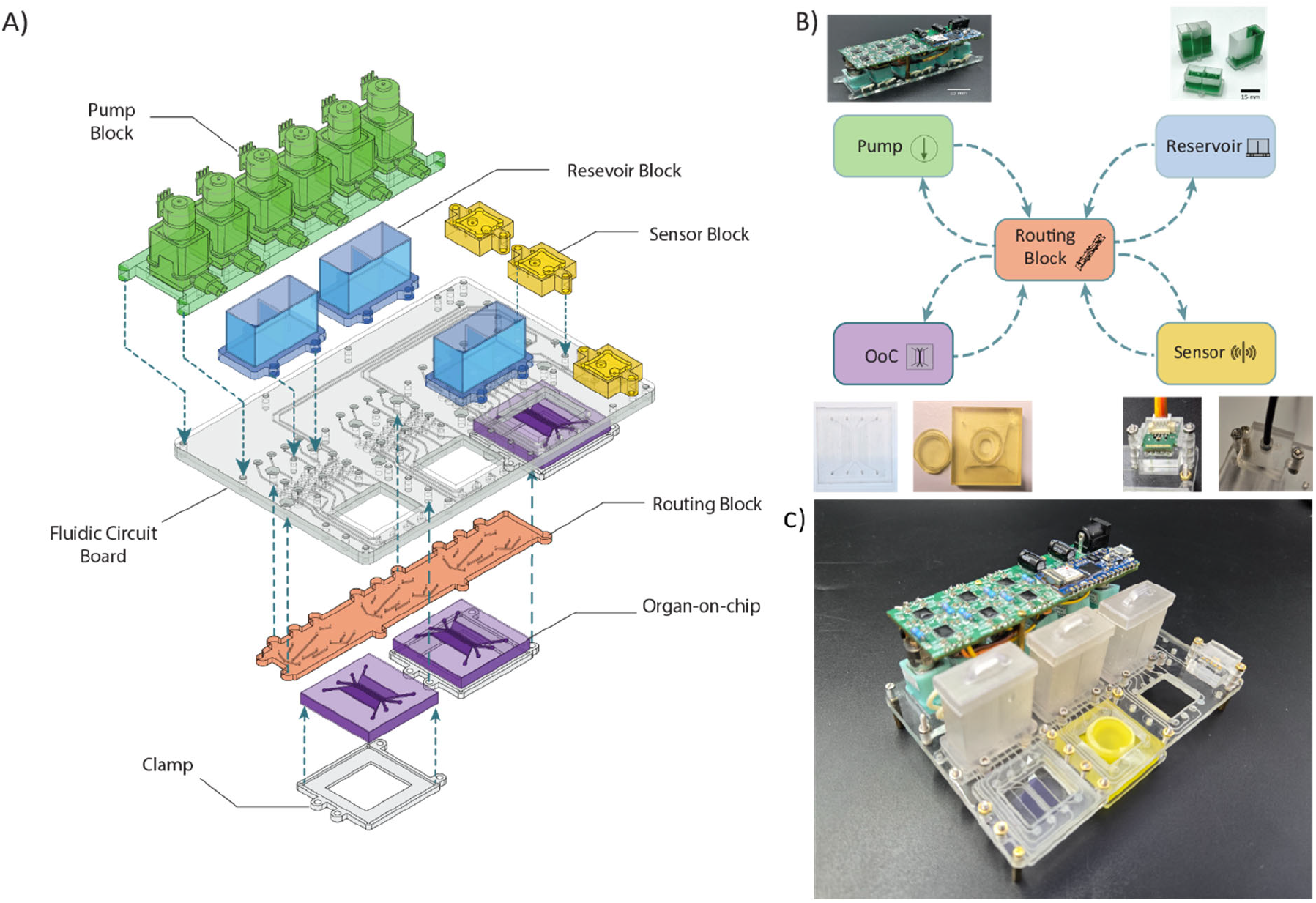
a) Exploded schematic view of the platform. Top side connections – pump block, reservoir block, sensor block ; Bottom side connections – routing block, organ-on-chips. b) Architectural schematic of the platform. All modules have inlet/outlet connections to the routing block. c) Fully assembled STARTER

The architecture opens up the possibility to utilize various MFBBs in custom configurations. In this work, the selection of MFBBs is catered towards a generic OoC experiment and aims to showcase the types of MFBBs that can be integrated into STARTER for such applications. Furthermore, this particular set of MFBBs can combine within the footprint of the FCB to enable fully automated and standalone experiments with only a 12 Volt power adaptor as a peripheral. The concept of being-stand-alone is not limited to these specific MFBBs however and can be implemented with other MFBBs with similar functionalities.

We also publish detailed designs of our MFBBs and FCB to provide insight for other developers to design OoC technology with ISO 22916 compatibility in mind (Figure S1). ISO 22916 specifies guidelines regarding the external footprint and fluidic port locations of an MFBB. In particular, this ISO allows port positions to be specified on a 1.5 mm grid and the footprint dimensions to be specified in increments of 15 mm^[4]^. Therefore, ISO 22916 allows a wide range of port and footprint selections, offering freedom to module developers while imposing common-sense constraints. In this work, we further streamline our design process by following the TOP Design Rules (TDRs), a publicly available set of ISO 22916 compliant footprints and port locations. This set is available online and all the MFBBs shown in this work are compliant with the TDRs^[32]^. Table 1 summarizes the MFBBs utilized in this work.

**Table 1.**
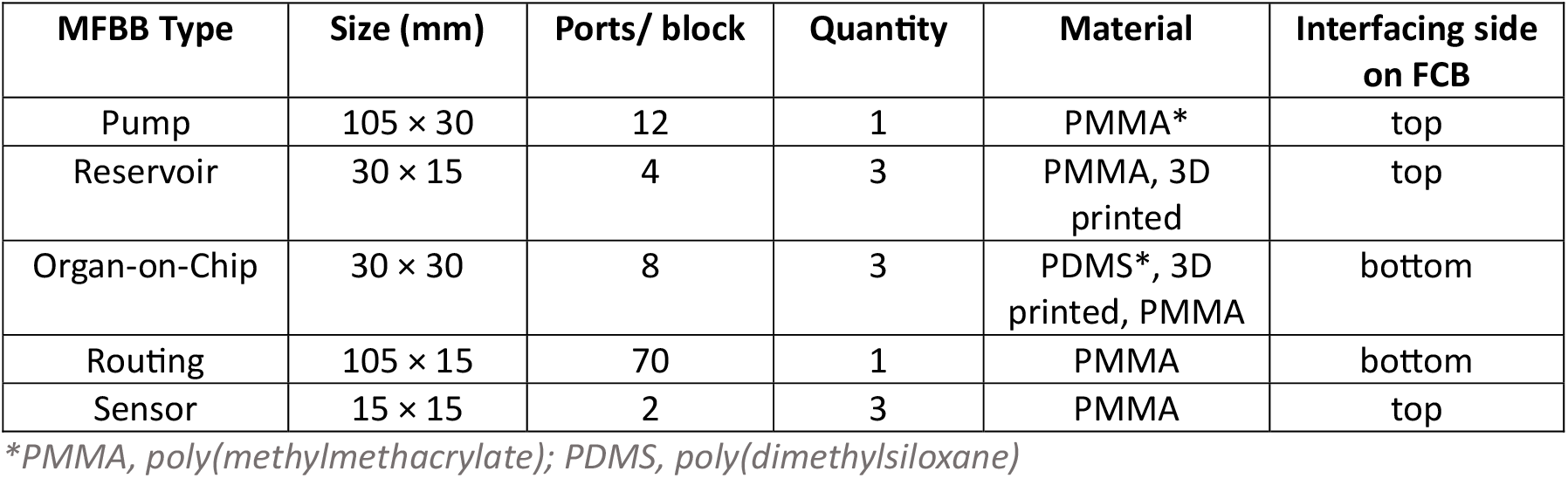
– Microfluidic Building Blocks in STARTER

| MFBB Type | Size (mm) | Ports/ block | Quantity | Material | Interfacing side on FCB |
| --- | --- | --- | --- | --- | --- |
| Pump | 105 × 30 | 12 | 1 | PMMA* | top |
| Reservoir | 30 × 15 | 4 | 3 | PMMA, 3D printed | top |
| Organ-on-Chip | 30 × 30 | 8 | 3 | PDMS*, 3D printed, PMMA | bottom |
| Routing | 105 × 15 | 70 | 1 | PMMA | bottom |
| Sensor | 15 × 15 | 2 | 3 | PMMA | top |

\*PMMA, poly(methylmethacrylate); PDMS, poly(dimethylsiloxane)

### 2.1 Components

#### 2.1.1 Fluidic Circuit Board

The FCB has a footprint of 127.75 mm × 85.5 mm, which makes it compatible with microtiter plate holders. MFBBs can be connected from the top or bottom side and are interconnected via 400 µm square channels in the FCB. All MFBBs connect to 400 µm diameter fluidic ports in the board connected to the channels in the FCB, with the locations of the ports on the FCB precisely matching those of the MFBBs. The layout is shown in Figure 1a, along with the schematic of the architecture in Figure 1b. Leak-free connections are made using O-rings recessed into the board that act as gaskets around the fluidic ports. The MFBBs are fastened onto the board using screws and nuts. The board has cut-outs that function as imaging windows or access points for inserts, depending on the type of the OoC. The FCB is either made of poly(methylmethacrylate) (PMMA) or cyclic olefin copolymer (COC) depending on if it is made in-house for iteration purposes or fabrication is outsourced to third-party suppliers.

#### 2.1.2 Routing Block

All ports on the platform are fluidically connected via the FCB to a single module called the ‘routing block’. The routing block is located in the central region of the FCB and completes all fluidic circuits to dictate the fluidic routing between all MFBBs on the board. This enables complete reconfigurability of all the fluidic circuits on the platform by just replacing the routing block. The routing block has a 105 mm × 15 mm footprint and connects to the bottom of the FCB.

Figure 2 shows two examples of different circuit configurations realized on the platform and corresponding routing blocks. However, this reconfigurability is not limited to the examples shown here. Together, this FCB architecture and routing module offer the flexibility of realizing a highly customizable set of fluidic circuits on the platform.

**Figure 2.**
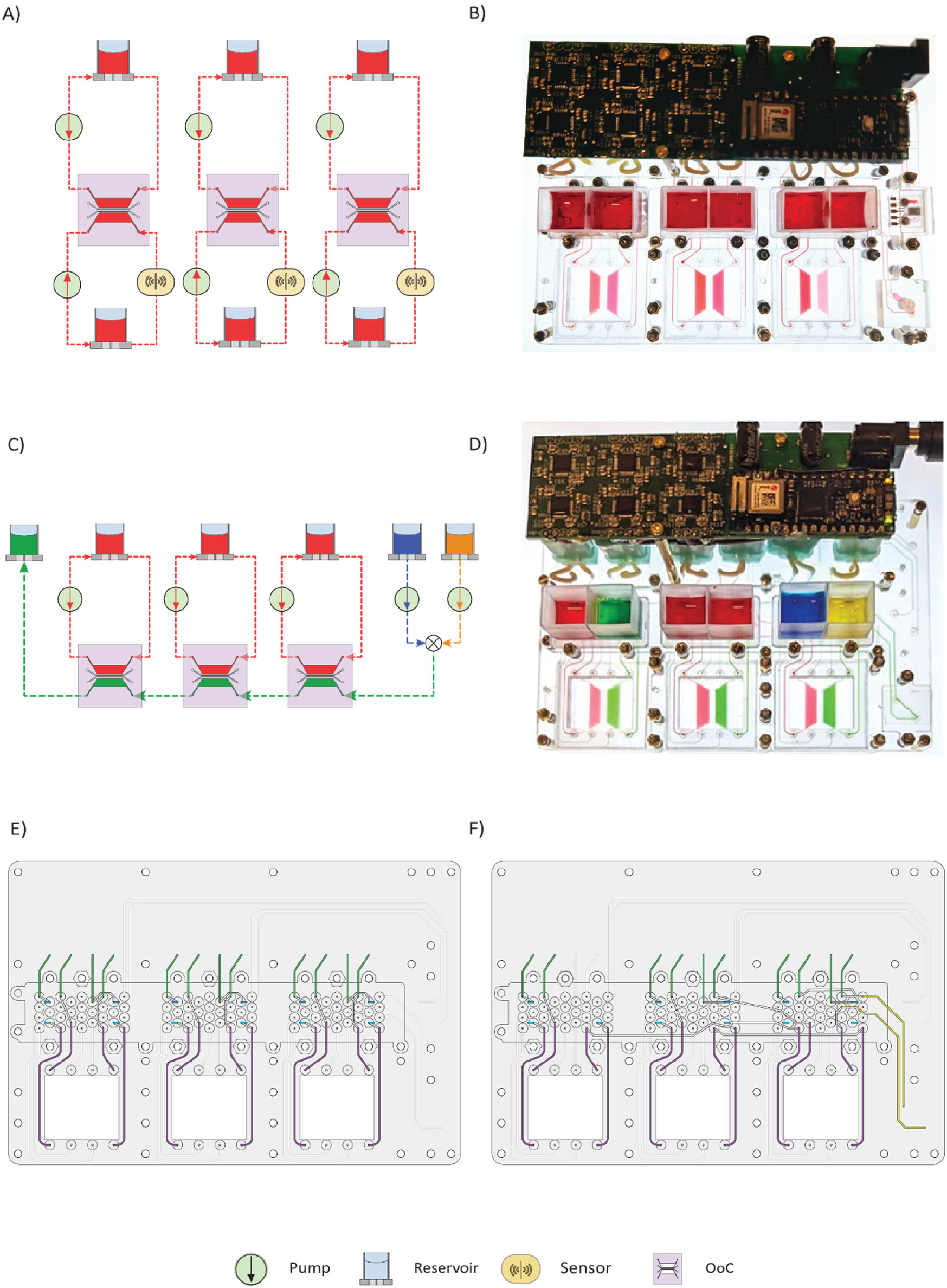
a) Schematic of Configuration 1 – all loops are running in isolation. Two channels per chip have pumping and one loop has an in-line sensor. b) Image of STARTER running Configuration 1. c) Schematic of Configuration 2 – One channel per chip is running in isolation. One channel per chip is in series after mixing of two reservoirs. d) Image of STARTER running Configuration 2. e) and f) show channel connection on the FCB for Configuration 1 and Configuration 2 respectively.

#### 2.1.3 Pump Block

This MFBB enables fluidic circulation on the platform. The pump block is comprised of six commercial peristaltic pumps driven by a custom printed circuit board (PCB) that houses an onboard Arduino. This enables wireless control of the pumps over low-energy Bluetooth (BLE). The electronics and pumps are integrated into a 105 mm × 30 mm block that connects with the FCB, powered by a single power cable. The pumps have a flow rate range of 0.18 µL/min to 180 µL/min using medical-grade PharMed tubing. The pumps and the tubes are housed in a PMMA baseplate that converts the entire block into an ISO-compliant MFBB. The tubes make a leak-free interface with the baseplate mounted to the FCB.

#### 2.1.4 Sensor Blocks

The FCB can integrate three 15 mm × 15 mm MFBBs, which can be chosen for in-line sensor blocks. This work shows a flow sensor and optical pH sensor module, both adapted from existing off-the-shelf products. The flow sensing module is built by making custom manifolds around a commercial thermal flow sensor chip. The pH module integrates a commercially available optical pH sensor spot into a microfluidic channel of a 15 mm × 15 mm MFBB. The pH module is designed so that an optical fiber aligns exactly above the location of the sensor spot.

#### 2.1.5 Various Organ-on-Chips

Compatible OoC MFBBs have a footprint of 30 mm × 30 mm and port location as specified by the TDRs. This FCB accommodates OoCs with 8 ports (4 inlets and 4 outlets) that can reversibly interface with the board via a clamp. Overviews of different OoC materials have been reported along with their specific applications ^[29]^. Yet, no platform prior to this work has demonstrated interfacing with OoCs of different make, organ types, and materials. Here, we first use PDMS chips closed with coverslips as example OoC devices. This particular PDMS OoC has two wide outer channels and two narrow channels used for endothelial cell culture to form a VoC model (Figure 4d). The broader outer channels are designed to act as ‘worst case’ design in terms of bubble formation and accumulation and can be used to technically monitor bubbles during extended operations. In addition to the PDMS chips, previously reported Intestinal Explant Barrier Chips (IEBC)^[33]^ were adapted according to the TDRs to conform to a 30 mm × 30 mm MFBB footprint. This enabled the integration of the 3D-printed OoCs onto the platform via the same clamps used for the PDMS devices. However, the types of OoCs compatible with the platform are not limited to these two particular examples, as the strategy of clamping OoCs onto the FCB is agnostic to the OoC material and architecture.

**Figure 3.**
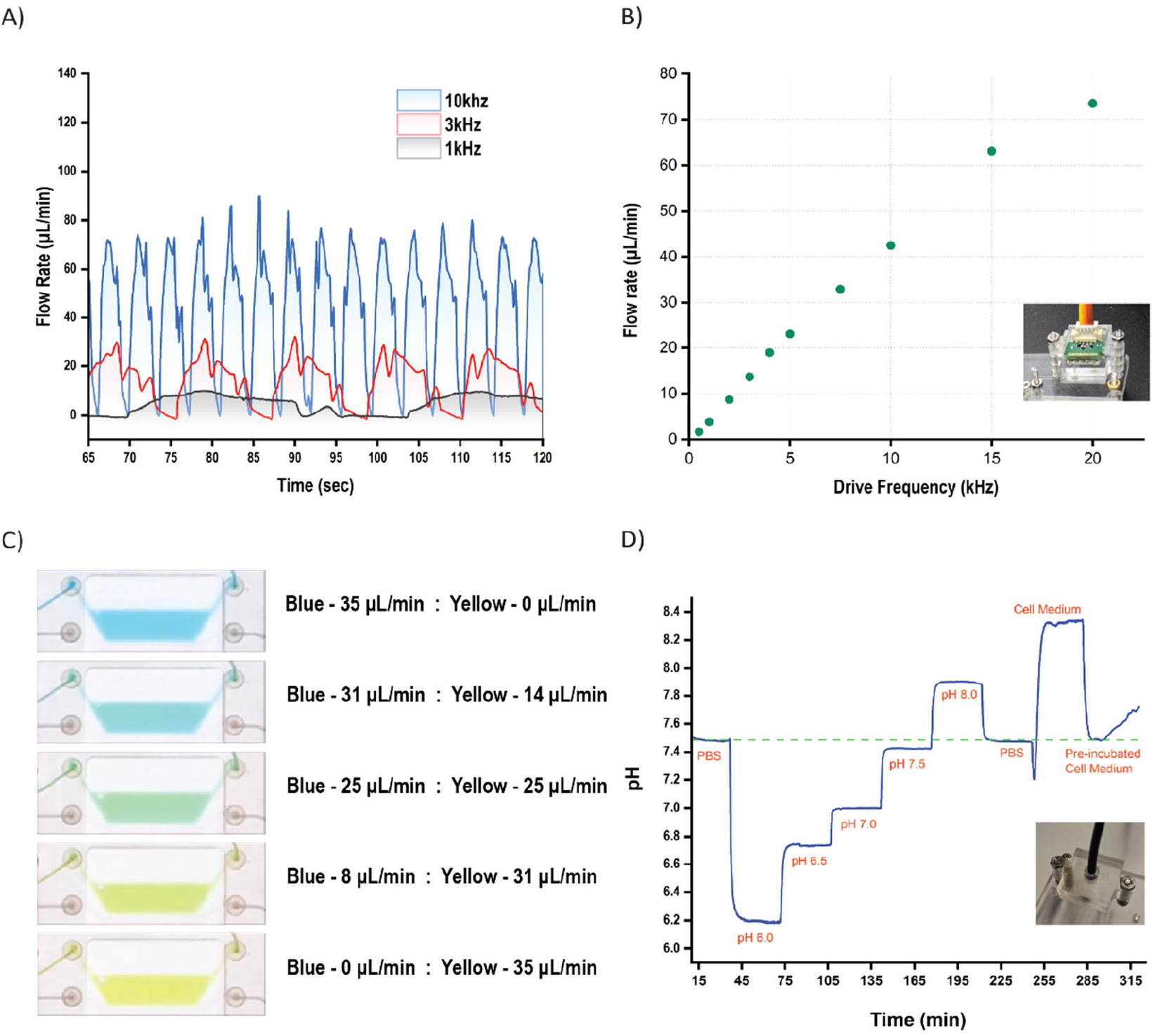
a) Flow profiles at different frequencies. b) Average flow rates for various input pump frequencies. The inset is a picture of the flow sensing module. c) Color gradient by varying dosing from blue and yellow reservoirs. d) pH measurements and characterization for varying pH solutions with an inset of the optical sensing module.

**Figure 4.**
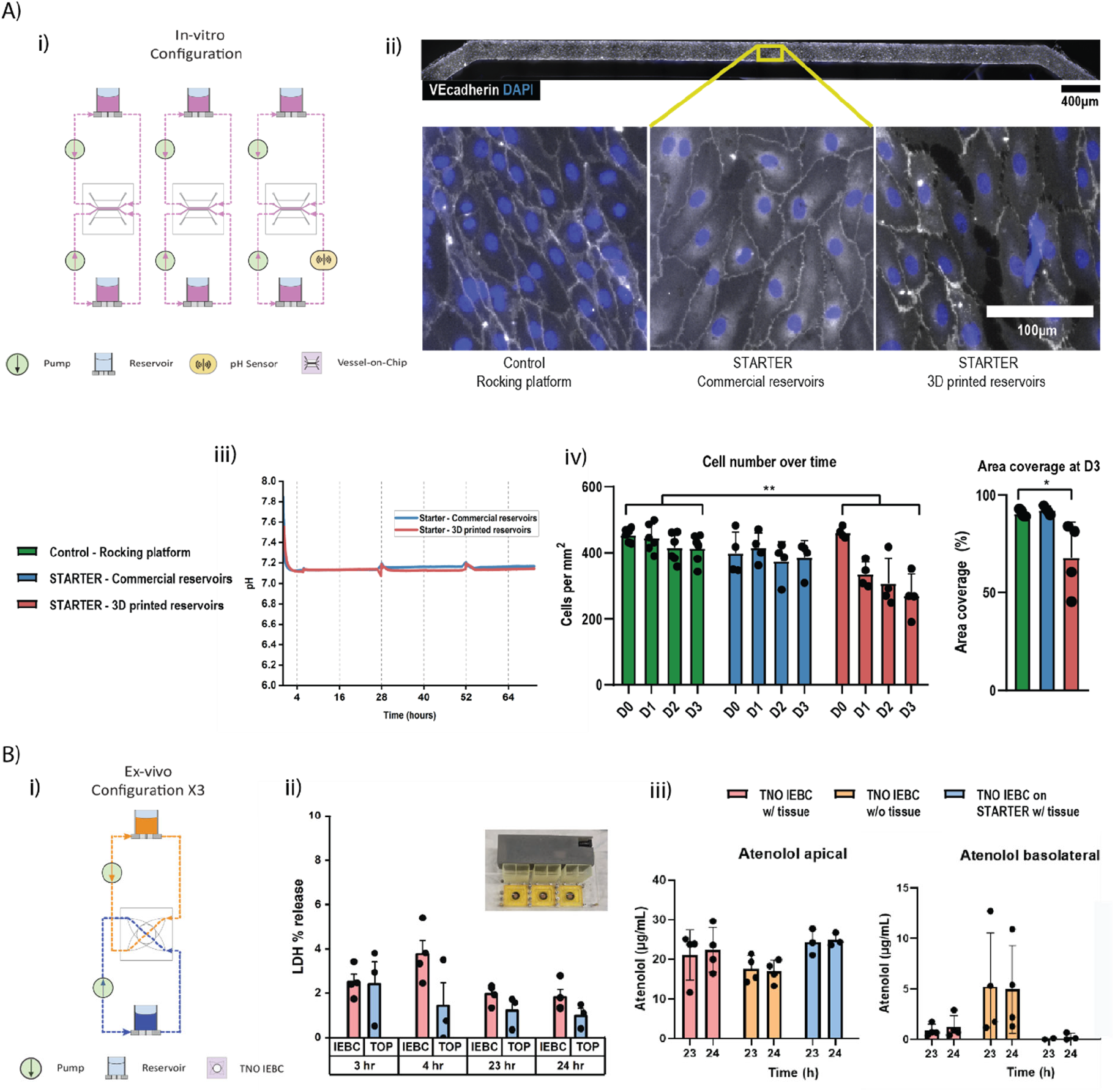
a) i) Schematic of the configuration for the in-vitro experiments. ii) Fluorescence Microscopy images of VoC channels for 3 conditions. iii) pH measurement during the in-vitro experiment. Iv) Cell Number and cell coverage analysis comparing control, STARTER with two types of reservoirs. b) i) Schematic of the configuration for the ex-vivo experiments. This loop was repeated three times on STARTER. ii) LDH secretion comparison between IEBC on STARTER and IEBC controls. iii) Atenolol transport in apical and basolateral sides. *(P<0.05), **(p<0.005)

#### 2.1.6 Reservoir Blocks

The reservoir block stores the liquid that circulates between the OoCs, pumps, and the sensors. The platform allows the attachment of 3 reservoir MFBBs at once, each having a footprint of 30 mm × 15 mm. Each block has 4 ports distributed between two reservoirs per MFBB. This, therefore, amounts to 6 reservoirs on the platform. The reservoirs can have varying designs based on volume requirements and customizations. In this work, we showcase 3D-printed reservoirs as well as ISO 22916 compliant MFBBs built around commercially available reservoirs. (SI)

## 3 Results

### 3.1 Fluidic Interfacing

Low durometer silicone O-rings are used to form a reliable seal while interfacing both soft and hard materials. This is also a differentiating factor compared to previously reported TOP platforms that used Vitron O-rings^[28]^. Our choice of O-rings makes this platform universal to various MFBB materials, as shown in Table 1. The screw arrangements are designed so that each MFBB can be detached from the FCB independent of the other MFBBs. This is especially useful in preparing the platform for perfusion experiments, where the priming of the platform and the initial culture of the OoC can be done independently. (Figure S3) The cultured OoCs can be swapped on and off the platform without generating bubbles at the fluidic ports. This is possible due to the hydrophobicity of the chosen O-rings which allows the port to retain a hanging drop when an OoC is removed. The hanging drop enables liquid-liquid connection when reconnecting the already filled OoC, allowing bubble-free swapping of the OoC.

### 3.2 Microfluidic Operation

As previously mentioned, the fluidic circuits of STARTER can be reconfigured by replacing the routing block and to demonstrate this, we show two specific configurations along with the fluidic routing. (Figure 2)

#### 3.2.1 Parallel Operation

As depicted in Figure 2a, this fluidic circuit configuration enables the parallel operation of 3 identical OoC experiments with the perfusion of two channels per chip. The OoCs used are made of PDMS and bonded with coverslips (as described in section 2.5), each OoC consisting of 4 individual channels corresponding to 4 inlets and 4 outlets. The pumps are distributed to perfuse two channels per chip to perform identical parallel and isolated operations of the 3 OoCs. Two channels of each OoC are connected to a pump and reservoir, and one channel out of these two channels has the possibility of an in-line sensor. The experimental validation is shown in Figure 2b, where red food dye is circulated in isolated loops at a flow rate of 27 µL/min. A variation of this configuration was used to perform cell culture experiments.

#### 3.2.2 Combined Operation

Figure 2c depicts the fluidic circuit that combines parallel and series operation within the 3 OoCs. The OoCs, reservoirs, and pump block are the same as described before, with the only change being a different routing block. In this fluidic circuit, each OoC has one channel under isolated circulation (red) and one in combined circulation (green). For the series circuit, liquids from two reservoirs (blue and yellow) get mixed in the routing block before flowing into the inlet of the first OoC (right) through a MFBB that serves as a non-functional substitute for a sensor. The outlet of this channel from the first OoC is connected to the inlet of the next OoC channel (middle), which similarly connects to the inlet of the third OoC (left). This is visible as the green color being perfused through the channels connected in series (right to left in Figure 2d). This configuration utilizes 5 pumps and 6 reservoirs and is visualized using the respective food dyes as depicted in the schematic. The flow rate applied is 27 µL/min, and the mixing flow rate variations are shown in Figure 3c.

### 3.3 Sensor measurements

The platform has the capability to integrate three 15 mm × 15 mm sensor modules. Here, we demonstrate two sensors that can be applied in an OoC experiment. First, a commercial thermal flow sensor is converted into a module and used to characterize the pump block. It is also used to measure the flow rate variations over time for different flow rates. In addition to the flow sensor, a commercial optical fiber based sensor is integrated to showcase the capability of utilizing optical in-line sensors. In this case, a pH sensor is integrated and characterized before applying it during a cell culture experiment.

#### 3.3.1 Flow Sensing

The pumps in the pump block are characterized over a flow range of 0.5-120 µL/min, as this was the measurement range of the selected flow sensor. The sensor measures the flow rate variation for a particular pump frequency, as shown in Figure 3a. This flow variation is then averaged to obtain average flow rates at specific frequencies. Figure 3b shows the measured average flow rates with respect to specific operating frequencies. An almost linear relation between the pump frequency and flow rate was observed. The monitoring of flow variation can be particularly useful for experiments where change in flow rates is critical. Furthermore, by running two peristaltic pumps with each other, we show mixing capabilities and also obtain a concentration gradient. This is demonstrated by mixing blue and yellow colors to obtain different shades of green by varying the ratio of the respective flow rates. (Figure 3c). We also compare the flow rate variation of two pumps running together against the isolated pump performance. (Figure S2)

#### 3.3.2 pH Sensing

Optical pH sensor spots that had a range of pH 6.0 - pH 8.0 were chosen, considering the physiological samples used in OoCs. Solutions of varying pH were measured to characterize the sensors, resulting in step curves, as seen in Figure 3c. PBS was used as a reference to check for drift before and after pH solutions varying from 6.0 to 8.0 in steps of 0.5 were flowed through at a flow rate of 27 µL/min. pH of room temperature and pre-incubated cell medium was also measured. The pH sensor readout was observed to be more accurate around pH 7.0 and had a maximum deviation of 0.2 pH at pH 6.2. No drift was observed over 3 hours. Finally, these pH sensor spots were used to monitor the pH on two platforms over a 3-day cell culture.

### 3.4 Biological Experiments

#### 3.4.1 Vessel-on-chip perfusion

The platform was tested for compatibility with automated cell culture experiments. A VoC was selected as an example OoC system as they are known to benefit from recirculating unidirectional medium flow. For this experiment, the middle channels of the previously described OoCs were perfused in isolated loops (Figure 4a i). Human umbilical vein endothelial cells (HUVECs) cultured under continuous flow conditions of 1 µL/min for 3 days on the platform showed characteristic cobblestone morphology, a cortical filamentous actin pattern, and expressed endothelial cell marker vascular endothelial (VE)-cadherin, indicative of a healthy endothelium (Figure 4a ii). Two platforms were used during the experiment – one with 3D-printed reservoirs and one with commercially available reservoirs. It was observed that the cells performed better in terms of cell coverage with commercial reservoirs compared to the 3D-printed reservoirs (Figure 4 a iv). This difference is suspected to be due to leaching of the 3D printed resins on exposure to daily fluorescence imaging but is comparable to additional controls with pro-inflammatory stimulus (Figure S4). Furthermore, the OoCs on the platform with commercial reservoirs were identical to control OoCs that were not connected to the FCB and under slow bidirectional perfusion on a rocking platform. No significant pH change was measured over the course of the cell experiments, with both platforms showing pH 7.1 (Figure 4a iii). This demonstrates the platform’s suitability for microfluidic OoC experiments traditionally perfused with rocking platforms or external pumps.

#### 3.4.2 Intestinal explant barrier chip (gut-on-chip) culture

To further showcase the versatility and translatability of the platform, IEBCs were integrated into the platform. The IEBC allows perfusion on both sides of a scaffold with intestinal tissue, thus enabling the possibility of conducting gut permeability assays. Freshly obtained porcine gut explants were inserted into the chips after the channels were primed using the platform. Flow rates of 33 µL/min were applied at either side of the explant tissue for 24 hours. The reservoirs were sampled at fixed intervals (3, 4, 23, and 24 hours) and checked for lactate dehydrogenase (LDH) secretion from the tissues to determine cellular viability. The concentration of LDH produced by IEBCs on the platform was then compared to control IEBCs perfused via the standard setup^[33]^. The results confirmed proper tissue viability in both setups with a similar profile over the entire course of the experiment.

Furthermore, tissue barrier function was confirmed in both setups by measuring the transport of atenolol, a low permeability compound that translocates across the intestinal barrier paracellularly. Results showed that transport in both setups was again comparable, with low transport of atenolol across the intestinal barrier, confirming intact tight junctions. Notably, transport in both setups was significantly lower than that in control IEBCs without tissue included. We, therefore, showcase comparable performances of established IEBCs on the platform, which features a much more compact footprint (Figure S5) and offers possibilities for future sensor integration and customization.

## 4 Conclusion and Outlook

In this work we introduce STARTER, a stand-alone, modular reconfigurable platform based on TOP and demonstrate its versatility for designing and carrying out OoC experiments. STARTER accommodates both tissue culture insert-based and microfluidic channel-based OoC models, agnostic to the substrate material. Pumps, reservoirs and sensors are all integrated within an ANSI/SLAS microplate footprint enabling dynamic monitoring of automated multi-OoC experiments in a compact portable package. The fluidic circuits can be customized as per requirements, thus offering experimental freedom with the integrated modules. Demonstrations of mixing and metering, pump characterization and sensor characterization highlight the technical capabilities of our integrated system. To highlight the applicability of STARTER’s versatility in OoC experiments, both in-vitro and ex-vivo studies were performed. As an exemplar in-vitro experiment, HUVECs were cultured over the course of 3 days in a vessel-on-chip model with continuous pH monitoring. The results confirm similar cell numbers and coverage to controls while revealing difference between 3D-printed and commercially available reservoirs. Furthermore, ex-vivo experiments were conducted in previously reported OoCs on STARTER over 24 hours. Cell viability and barrier function of a porcine gut explant was assessed and shown to be comparable to controls in the traditional setup. These results showcase the advantage of STARTER in reducing experimental footprint, adding functionality and versatile integration without compromising on biological performance.

The design of STARTER is complaint with ISO 22916 and specifically the TOP Design rules (TDRs) which is a specific implementation of ISO complaint footprints and port layouts. A wide variety of modules become eligible for integration as long as the port locations and footprints adhere to these standardized design guidelines. In this work, we showcase a few possible module combinations along with a general strategy for the implementation of the standardized designs in modular microfluidic systems. The list of components used in this work as well as design files are all made freely available in an open source environment - <u>**GitHub - TOP-OoC/Starter-Kit**</u>. These openly accessible resources will serve as a ‘STARTER kit’ for easier adoption and implementation by developers at this nascent stage of standardizing modules. Additional resources such as ISO explainer documents, TDR guidelines and an automated fluidic routing tool (<u>**MMFT Routing Block Channel Router**</u>) are also available. The fluidic routing tool is developed by collaborators in Technical University of Munich (TUM) with renowned experience in design automation^[34]^ and is especially useful in designing routing blocks for customized applications. These resources aim to lower the threshold of adoption of standardized designs, particularly in the field of OoC.

The architecture of STARTER can serve as a foundation for generic platforms aiming to add perfusion to existing modules in a portable footprint. The ability to have a standalone and reuseable platform makes it truly translational. However, implementing STARTER in external lab settings is a key next step in validating the translatability, design choices, interfacing and robustness of the platform. A natural progression for wider adoption would involve simplifying the mechanical connections to improve useability. This will particularly benefit time constraining and space restricted workflows. Future work on reducing electrical connectors is also of vital importance. As the number of integrated sensors increase in a compact form factor, the web of cables running to these sensors can cause congestion and hamper useability. A method to further integrate the electronics and sensor read-out on the platform will enhance ease-of-use while benefiting from an already matured electronics industry. Integration of valves in the routing block would enable active re-programmability of the platform compared to the current method of replacing the block itself. This will allow further possibilities of multiplexing, pumping and fluidic operations on the platform.

Dissemination of the benefits of standardized designs also becomes vital to ignite adoption by component developers. The ISO 22916 standard could harmonize international academic and commercial entities, initiating an interoperable market. The benefits of the ISO standard is not limited to module developers alone, the growing ecosystem of complaint components can be used by system designers to realize custom platforms for their specific applications. In a wider perspective, a generic open-source platform tackles fundamental obstacles in the industrial adoption of MPS systems, as emphasized by various global committees.^[35]^ Standardized design principles enable the development of tailored OoC models and platforms with integrated automation, facilitating rapid iteration of more novel and complex models. An OoC model developed for a standardized platform such as STARTER has the potential to be adapted for high-throughput systems that follow the same standards in the future ^[36]^. This translational capabilities not only strengthens collaboration but also supports continuous improvements informed by user feedback and preferences. This approach can bridge the gap between academia and the pharmaceutical industry by first validating experimental models on generic platforms, then enabling their scaled automation once validated. Therefore, embracing an open-source approach fosters collaborative development of solutions aimed at enhancing robustness, reproducibility, and technical maturity, ultimately facilitating future regulatory approval processes.

In conclusion, we introduce a novel generic platform designed for stand-alone multi-OoC experiments, with in-line sensing and fluidic reconfigurability capabilities. The versatility of the architecture combined with a reversible and material agnostic integration strategy makes STARTER suitable for a wide range of applications. To the best of our knowledge, this represents the first application of standardized designs on a universal platform capable of accommodating diverse OoC models from multiple suppliers. The applications extend beyond OoC experiments demonstrated in this work and can include module testing, quality control, benchmarking, integration tests, and automated microfluidics, all within a portable form factor. The open access dissemination of resources will foster broader collaboration and contribution of new designs from the community. Eventually, we expect that development, testing and implementation of new OoC applications will be strongly accelerated, both in the setting of early R&D and in commercial product development. The paradigm of open-source design represents a significant breakthrough in efficiency and innovation by creating a precompetitive domain in a field that has been converging towards ‘point solutions’ and proprietary platforms. The boom in development, validation and testing of OoCs will, in-turn stride towards the ultimate objective of wider adoption of micro-physiological systems.

## 5 Materials and Methods

### 5.1 Assembly

The MFFBs and clamps were connected to the FCB using M2 nuts and bolts. 40° shore-A silicone O-rings (Gteek) with dimensions 1.02 mm × 0.74 mm were recessed at the locations of the fluidic ports. The O-rings were manually placed into the recesses in the FCB before connecting the MFBBs with screws onto the FCB. The compression of the soft O-rings with the screws enabled leak-free integration. Sufficient O-ring compression was verified for O-ring pocket depths varying from 0.8 µm to 0.9 µm and a pocket diameter of 1.35 mm.

### 5.2 FCB, MFBBs and Clamps

The OoCs were placed inside the clamp with ports facing toward the FCB. The clamp was then screwed onto the FCB similar to other MFBBs The FCB, MFBBs and clamps were designed in SolidWorks© (2022). The FCB was made of COC and manufactured by Micronit B.V. (The Netherlands). The clamps, and auxiliary MFBB parts were made by micromilling (Datron Neo, Germany) PMMA. The routing block was micromilled similarly and closed off with medical grade double sided pressure sensitive adhesive tape (PSA, ARcare 90445Q). The bases for the pump block, sensor blocks and commercial reservoirs were manufactured similarly out of PMMA. It is also possible to fabricate the FCB in-house by micromilling PMMA and bonding with PSA tape as described above. All designs are shown in SI.

### 5.3 Organ-on-Chips

PDMS (Sylgard 184 elastomer kit) was mixed (10:1 base polymer to curing agent (w/w)) and then degassed for 1 h. Degassed PDMS was then cast into a micromilled double sided PMMA mold and degassed again for 1h. The baseplate provides the features for the fluidic pathing as well as cutting lines for 6 devices, while the top controls chip thickness and levelness. The degassed mold was placed at 60 °C for at least 3 h. The mold was then disassembled, and the individual microfluidic devices were cut out according to the cutting guides, resulting in uniform 30×30 mm PDMS slabs. Subsequently, inlets and outlets were punched out using 1 mm biopsy punch (Ted Pella, Inc., USA). A clamp orients a single PDMS slab while a second micromilled punching guide was aligned to the PDMS using M2 bolts as guide pins. Inlet and outlet locations were punched according to the TDRs and the PDMS slabs were plasma bonded to 24×24 mm coverslips (VWR). The final devices were placed at 60 °C for at least 1 h.

### 5.4 Pump Block

This module uses six stepper-motor driven peristaltic pumps, manufactured by Takasago Fluidic Systems. The stepper motors have a rated pump rate of 0.18 -180 µL/min. A custom pump driver PCB was used which was controlled by an Arduino Nano 33 IoT. Six unpopulated spaces for through-hole resistors were left in the PCB for pump-specific feedback resistors to be added. These resistors control the current output by the driver and must be added during or after manufacturing. For the pumps listed here, a resistor value of 56 kOhm was used. A custom firmware for the pump driver PCB allowed individual control of the pumps via BLE.

The pumps were attached to a micromilled PMMA base plate that connected the array of pumps to the FCB. The PCB was connected to the Pump Block Base with M2 PCB standoffs. The PCB was conformal coated with epoxy before use in high humidity settings like incubators. All design files are available in the previously mentioned Github page.

### 5.5 Reservoirs

The reservoirs were either 3D printed or converted using commercially available components (4.5mL Interaction Tanks Microfluidic ChipShop). For the 3D printed reservoirs, designs were created in SolidWorks© 2022 and sliced in PreForm slicing software; Formlabs BioMed Clear v1.0 resin was used in a FormLabs 3B+ printer. The printed reservoirs were then washed in 100% IPA under agitation (FormLab Form Wash) followed by a 2-hour UV cure (Form Cure) and a final overnight bake at 80 °C. The bottom of the reservoirs (side interfacing with the FCB) was sanded to obtain a flat surface both in terms of roughness and curvature. In case of the commercial reservoirs, a 30mm x 15mm base plate with required ports were milled onto which the reservoirs were attached by a press fit.

### 5.6 Flow Sensor measurements

The flow sensor used was the LPG10-1000 (Sensirion AG) and logged in the Sensirion Viewer software. The data was logged as .csv files and plotted later using OriginPro (2024). The flow rate was measured at different pump frequencies for a duration of 5 min for each data point. The average flow rate was then taken over the 5 min duration.

### 5.7 pH measurements

The pH sensor spot used was a commercial product (PHSP5-PK6) from PyroScience GmBH and the measurements were conducted using SPFIB-BARE optical fibers connected to a Firesting Pro. Prior to the measurements, a 2-point calibration was performed as suggested by the supplier using the recommended pH 2 (PHCAL2) and pH 11 (PHCAL11) calibration capsules. The calibration was done in ambient conditions under a flow rate of 15 µL/min. For the pH measurements, buffer solutions of pH levels ranging from 6.0-8.0 were prepared in posphate buffered saline (PBS) using 0.1 M NaOH and flowed through the platform. Each buffer solution was allowed to recirculate for 30 minutes followed by a dry air run of 10 min to ensure complete removal of the previous solution prior to introducing the next solution.

### 5.8 Vessel-on-Chip Cell Seeding

Green fluorescent protein-tagged human umbilical vein endothelial cells (GFP-HUVECs, ZHC-2402, Cellworks) were cultured as suggested by the supplier. In short, cells were expanded on 0.1 mg/ml collagen-1 (rat tail collagen-1, Gibco) coated T75 flasks in endothelial cell growth medium-2 (ECGM-2, PromoCell) supplemented with penicillin-streptomycin (50 U/mL, Gibco) at 37 °C in humidified air with 5% CO_2_. To facilitate cell adhesion in the Vessel-on-Chip (VoC) channels of the PDMS chip, PDMS surface was functionalized with 2 mg/mL dopamine (Sigma-Aldrich) in 10 mM Tris-HCl buffer (pH 8.5) for 1 hour at room temperature (RT), followed by 3 washes with sterile filtered deionized water and finalized with a 0.1 mg/mL collagen-1 coating for 30 minutes at 37 °C. Afterwards, the channels were washed with ECGM-2 to remove non-bound collagen. HUVECs were then obtained from confluent flasks using 0.05% trypsin-EDTA (Gibco), seeded at 2·10^6^ cells/mL and incubated for 1 hour at 37 °C followed by a wash of fresh medium to remove non-adhered cells. To enable full attachment of the cells to the VoC channel walls, the chips were kept static in 37 °C for 4 hours. Afterwards, VoCs were transferred onto a rocking platform providing bi-directional flow to ensure frequent medium refreshment (OrganoFlow, Mimetas, set at 10°, 1-hour intervals). Medium was refreshed twice per day and cells were allowed to form a monolayer prior to start of the experiment on STARTER.

### 5.9 Vessel-on-chip Analysis

The number of cells was monitored over the course of the 3-day experiment and compared between the VoCs connected to Starter with either commercial or 3D-printed reservoirs, as well as VoCs kept with bi-directional flow on a rocking platform in plain ECGM-2 or supplemented with pro-inflammatory cytokine TNF-α (2ng/ml) as positive and negative control respectively. For this, the GFP-tagged HUVECs were imaged daily starting directly after connecting the VoCs to Starter using the EVOS M5000 Imaging System. Cell numbers were determined in CellProfiler (version 4.2.8), in which individual cells were segmented using three-class Otsu thresholding method based on the GFP-intensity images. VoC channels were excluded from analysis if the amount of cells at the start of the experiment was less than 75% of the ECGM-2 conditions, or if technical faults resulted in sudden loss of cells.

Additionally, cell morphology was assessed using immunostaining of endothelial marker vascular endothelial-cadherin (VE-cadherin), cytoskeleton and nuclei. Directly after completion of the experiment, VoCs were removed from the Starterkit and HUVECs were fixed in 4% paraformaldehyde in PBS containing Ca^2+^ and Mg^2+^ for 10 minutes at RT. Afterwards, cells were permeabilized and blocked in permeabilization and blocking buffer (PBB) containing 0.1% Triton X-100 (Sigma-Aldrich) and 10 mg/mL bovine albumin serum (BSA, Sigma-Aldrich) in PBS for 60 minutes at RT. Afterwards, HUVECs were incubated with 5 µg/mL goat anti-human VE-cadherin (R&D systems) in PBB overnight at 4 °C. Extensive washing was performed to remove primary antibodies using 3 rinses and 3 10-minute incubations with PBS. Afterwards, HUVECs were incubated with 10 µg/mL donkey anti-goat Alexa Fluor 546, 4 droplets/mL ActinGreen (ThermoFischer Scientific) and 12.5 µg/mL 4’,6-diamidino-2-phenylindole (DAPI) in PBB for 4 hours at RT. After another set of extensive washing, samples were imaged using the EVOS FL Auto 2 Imaging System. VE-cadherin pattern was used as input in CellProfiler to determine the percentage coverage of the HUVECs in the VoC channel. Percentage cell coverage was determined in CellProfiler, in which the total area of the sum of the individual cells was determined with respect to the channel area. For this, cells were segmented first using three-class Otsu thresholding of the DAPI images, followed by two-class Otsu thresholding of the VE-cadherin images.

### 5.10 IEBC tissue culture

The STARTER platform and Explant Barrier Chips were prepared one day prior to an experiment. After assembling STARTER, reservoirs were filled and the system was subsequently flushed with 20% biofilm (Umweltanalytik) and afterwards flushed with PBS. Next, Williams E supplemented with 1% BSA was added for overnight incubation in a humidified incubator at 37 °C with 5% CO_2_. Flow rates of 33 µL/min were used for overnight incubation and flushing, respectively. The next day, systems were transferred to a working bench.

Porcine intestinal colon tissue from a healthy adult pig was obtained from a local slaughterhouse. No ethical approval was needed for the collection of intestinal tissue from these animals as the tissue was redundant to the slaughter procedure. Procedures for handling and processing the tissue was according to previously published methods ^[28,37]^. In brief, intestinal tissue was collected within 15 minutes upon death of the animal and immediately flushed with ice cold supplemented Williams E buffer to remove fecal content. During transportation and preparation in the lab, the tissue was placed in ice cold supplemented Williams E buffer. At the laboratory, fattissue and the musculo-serosal layer of the mucosal layer was dissected off and round segments of 11.1 mm in diameter (area of 0.968 cm^2^) were punched. Mounting of the segments into the IEBC occurred within 4 hours after intestinal tissue collection. All experiments were performed in compliance with Dutch legislation on the use of redundant human (AVG, WMO) and slaughterhouse porcine tissue, and institutional guidelines on handling human and animal tissue regarding safety and security.

1 mm thick EPDM rubber rings (Eriks), intestinal tissue segments (mucosal side upwards) on a woven mesh of 170 μm in thickness and 50% open area (Nitex, Sefar) and a fixing insert were clicked in the snap fit mechanism, thereby separating the apical and basolateral compartments of the microfluidic chip. Subsequently, the Williams E supplemented with 1% BSA was replaced by the apical and basolateral media: Williams E supplemented with FD4 (50μM) and atenolol (10 μM) and Williams E supplemented with 4% BSA, respectively. Thereafter, the system was placed back in the incubator and perfused at 33 µL/min. Apical and basolateral samples were collected from the medium reservoirs at previously mentioned time points. At the end of the experiment, tissue segments were flushed with warm PBS and removed from the Explant Barrier Chips and collected for subsequent analyses. The whole STARTER platform, tubings, chips and reservoirs were flushed and washed with first with 20% biofilm and then with 70% ethanol.

### 5.11 IEBC Analysis

[^3^H]Atenolol was applied as reference marker for paracellular transport and mixed with non-radiolabeled atenolol, to obtain final nominal concentrations of 10 μM in the apical solution with an associated radioactivity of 10 kBq mL^−1^. Transport was measured by taking apical (100 μL) and basolateral (500 μL) samples at indicated timepoints. Radioactive labelled compounds were measured using the Tri-Carb 3100TR Liquid Scintillation counter (LSC, Perkin Elmer, Boston Massachusetts, United States) after adding scintillation liquid (Ultima Gold, Perkin Elmer Inc., Boston, Massachusetts, United States) to the apical and basolateral samples.

To assess the viability of the *ex vivo* intestinal segments, the cytosolic enzyme lactate dehydrogenase (LDH) was measured in the apical and basolateral supernatants of the two-compartmental model using an LDH kit (Sigma-Aldrich). Intracellular LDH levels in control tissue segments collected before incubations were measured with the same kit, after homogenizing the tissue segments in ice-cold Williams E buffer using a Potter–Elvehjem type Teflon pestle tissue grinder (Braun) for 5 min at 200 rpm. Excreted LDH levels were expressed as percentage leakage of the total intracellular LDH of these blanc intestinal tissue segments. Samples were analyzed using the BioTek Synergy HT microplate reader (BioTek Instruments Inc., Winooski, VT) with an excitation/emission wavelength of 490 nm and 520 nm.

### 5.12 Statistics

Data are provided as the mean ± standard deviation or standard error of the mean. Differences between 2 groups were analyzed using 2-tailed Student’s *t* test; 1-way ANOVA with Tukey’s or Dunnett’s *post hoc* analysis was used for comparisons of multiple groups. Statistical significance was considered at *p* < 0.05, and calculations and graphs were generated using GraphPad Prism 8.0 (GraphPad Software Inc.) and Origin Pro (2024).

## Author Contributions

A.P. and E.R.S contributed equally to this work. A.P., E.R.S., J.L.Z., M.O., and A.D.v.d.M. conceptualized the study. A.P. and E.R.S., designed and tested the platform. L.E.d.H conducted in-vitro experiments and analyzed the data. B.d.W. and H.E.A designed IEBC for the platform. E.v.d.S. and K.W. conducted ex-vivo experiments and analyzed data. A.D.v.d.M., M.O., J.L.Z., and A.R.V supervised the research.

## Acknowledgements

This work was supported by the NWO-TTW Perspective Programme of the Dutch Research Council (NWO, project number P19-03) as part of the SMART Organ-on-Chip consortium. We acknowledge Henk Waayer for designing the pump block PCB and Sandro Meucci for assistance in fabricating the FCBs at Micronit B.V. (The Netherlands).

## References

1 Huh, D., Matthews, B. D., Mammoto, A., Montoya-Zavala, M., Hsin, H. Y., & Ingber, D. E. (2010). Reconstituting Organ-Level lung functions on a chip. Science, 328(5986), 1662–1668. 10.1126/science.1188302

2 Huh, D., Hamilton, G. A., & Ingber, D. E. (2011). From 3d cell culture to organs-on-chips. Trends in Cell Biology, 21(12), 745–754. 10.1016/j.tcb.2011.09.005

3 Vollertsen, A. R., Vivas, A., van Meer, B., van den Berg, A., Odijk, M., & van der Meer, A. D. (2021). Facilitating implementation of organs-on-chips by open platform technology. Biomicrofluidics, 15(5). 10.1063/5.0063428

4 Piergiovanni, M., Leite, S. B., Corvi, R., & Whelan, M. (2021). Standardisation needs for organ on Chip Devices. Lab on a Chip, 21(15), 2857–2868. 10.1039/d1lc00241d

5 Zhang, B., & Radisic, M. (2017). Organ-on-a-chip devices advance to market. Lab on a Chip, 17(14), 2395–2420. 10.1039/c6lc01554a

6 Xiao, S., Coppeta, J. R., Rogers, H. B., Isenberg, B. C., Zhu, J., Olalekan, S. A., McKinnon, K. E., Dokic, D., Rashedi, A. S., Haisenleder, D. J., Malpani, S. S., Arnold-Murray, C. A., Chen, K., Jiang, M., Bai, L., Nguyen, C. T., Zhang, J., Laronda, M. M., Hope, T. J., … Woodruff, T. K. (2017). A microfluidic culture model of the human reproductive tract and 28-day menstrual cycle. Nature Communications, 8(1). 10.1038/ncomms14584

7 Novak, R., Ingram, M., Marquez, S., Das, D., Delahanty, A., Herland, A., Maoz, B. M., Jeanty, S. S., Somayaji, M. R., Burt, M., Calamari, E., Chalkiadaki, A., Cho, A., Choe, Y., Chou, D. B., Cronce, M., Dauth, S., Divic, T., Fernandez-Alcon, J., …Ingber, D. E. (2020). Robotic Fluidic coupling and interrogation of multiple vascularized organ chips. Nature Biomedical Engineering, 4(4), 407–420. 10.1038/s41551-019-0497-x

8 Bas-Cristóbal Menéndez, A., Du, Z., van den Bosch, T. P., Othman, A., Gaio, N., Silvestri, C., Quirós, W., Lin, H., Korevaar, S., Merino, A., Mulder, J., & Hoogduijn, M. J. (2022). Creating a kidney organoid-vasculature interaction model using a novel organ-on-chip system. Scientific Reports, 12(1). 10.1038/s41598-022-24945-5

9 Maschmeyer, I., Lorenz, A. K., Schimek, K., Hasenberg, T., Ramme, A. P., Hübner, J., Lindner, M., Drewell, C., Bauer, S., Thomas, A., Sambo, N. S., Sonntag, F., Lauster, R., & Marx, U. (2015). A four-organ-chip for interconnected long-term co-culture of human intestine, liver, skin and kidney equivalents. Lab on a Chip, 15(12), 2688–2699. 10.1039/c5lc00392j

10 McAleer, C. W., Long, C. J., Elbrecht, D., Sasserath, T., Bridges, L. R., Rumsey, J. W., Martin, C., Schnepper, M., Wang, Y., Schuler, F., Roth, A. B., Funk, C., Shuler, M. L., & Hickman, J. J. (2019). Multi-organ system for the evaluation of efficacy and off-target toxicity of anticancer therapeutics. Science Translational Medicine, 11(497). 10.1126/scitranslmed.aav1386

11 Ronaldson-Bouchard, K., Teles, D., Yeager, K., Tavakol, D. N., Zhao, Y., Chramiec, A., Tagore, S., Summers, M., Stylianos, S., Tamargo, M., Lee, B. M., Halligan, S. P., Abaci, E. H., Guo, Z., Jacków, J., Pappalardo, A., Shih, J., Soni, R. K., Sonar, S., … Vunjak-Novakovic, G. (2022). A multi-organ chip with matured tissue niches linked by vascular flow. Nature Biomedical Engineering, 6(4), 351–371. 10.1038/s41551-022-00882-6

12 Edington, C. D., Chen, W. L., Geishecker, E., Kassis, T., Soenksen, L. R., Bhushan, B. M., Freake, D., Kirschner, J., Maass, C., Tsamandouras, N., Valdez, J., Cook, C. D., Parent, T., Snyder, S., Yu, J., Suter, E., Shockley, M., Velazquez, J., Velazquez, J. J., …Griffith, L. G. (2018). Interconnected microphysiological systems for Quantitative Biology and Pharmacology Studies. Scientific Reports, 8(1). 10.1038/s41598-018-22749-0

13 Azizgolshani, H., Coppeta, J. R., Vedula, E. M., Marr, E. E., Cain, B. P., Luu, R. J., Lech, M. P., Kann, S. H., Mulhern, T. J., Tandon, V., Tan, K., Haroutunian, N. J., Keegan, P., Rogers, M., Gard, A. L., Baldwin, K. B., de Souza, J. C., Hoefler, B. C., Bale, S. S., … Charest, J. L. (2021). High-throughput organ-on-chip platform with integrated programmable fluid flow and real-time sensing for complex tissue models in drug development workflows. Lab on a Chip, 21(8), 1454–1474. 10.1039/d1lc00067e

14 Wevers, N. R., van Vught, R., Wilschut, K. J., Nicolas, A., Chiang, C., Lanz, H. L., Trietsch, S. J., Joore, J., & Vulto, P. (2016). High-throughput compound evaluation on 3D networks of neurons and glia in a microfluidic platform. Scientific Reports, 6(1). 10.1038/srep38856

15 Nicolas, A., Schavemaker, F., Kosim, K., Kurek, D., Haarmans, M., Bulst, M., Lee, K., Wegner, S., Hankemeier, T., Joore, J., Domansky, K., Lanz, H. L., Vulto, P., & Trietsch, S. J. (2021). High throughput transepithelial electrical resistance (TEER) measurements on perfused membrane-free epithelia. Lab on a Chip, 21(9), 1676–1685. 10.1039/d0lc00770f

16 Zhang, Y. S., Aleman, J., Shin, S. R., Kilic, T., Kim, D., Mousavi Shaegh, S. A., Massa, S., Riahi, R., Chae, S., Hu, N., Avci, H., Zhang, W., Silvestri, A., Sanati Nezhad, A., Manbohi, A., De Ferrari, F., Polini, A., Calzone, G., Shaikh, N., … Khademhosseini, A. (2017). Multisensor-integrated organs-on-chips platform for automated and continual in situ monitoring of organoid behaviors. Proceedings of the National Academy of Sciences, 114(12). 10.1073/pnas.1612906114

17 Fantuzzo, J. A., Robles, D. A., Mirabella, V. R., Hart, R. P., Pang, Z. P., & Zahn, J. D. (2020). Development of a high-throughput arrayed neural circuitry platform using human induced neurons for drug screening applications. Lab on a Chip, 20(6), 1140–1152. 10.1039/c9lc01179j

18 Ong, L. J., Ching, T., Chong, L. H., Arora, S., Li, H., Hashimoto, M., DasGupta, R., Yuen, P. K., & Toh, Y.-C. (2019). Self-aligning Tetris-like (tile) modular microfluidic platform for mimicking multi-organ interactions. Lab on a Chip, 19(13), 2178–2191. 10.1039/c9lc00160c

19 Carvalho, D. J., Kip, A. M., Tegel, A., Stich, M., Krause, C., Romitti, M., Branca, C., Verhoeven, B., Costagliola, S., Moroni, L., & Giselbrecht, S. (2024). A modular microfluidic organoid platform using lego-like bricks. Advanced Healthcare Materials, 13(13). 10.1002/adhm.202303444

20 Loskill, P., Marcus, S. G., Mathur, A., Reese, W. M., & Healy, K. E. (2015). MORGANO: A LEGO®-like Plug & Play System for Modular Multi-Organ-Chips. PLOS ONE, 10(10). 10.1371/journal.pone.0139587

21 Schuster, B., Junkin, M., Kashaf, S. S., Romero-Calvo, I., Kirby, K., Matthews, J., Weber, C. R., Rzhetsky, A., White, K. P., & Tay, S. (2020). Automated microfluidic platform for dynamic and combinatorial drug screening of tumor organoids. Nature Communications, 11(1). 10.1038/s41467-020-19058-4

22 Schneider, S., Erdemann, F., Schneider, O., Hutschalik, T., & Loskill, P. (2020). Organ-on-a-disc: A platform technology for the centrifugal generation and culture of microphysiological 3D cell constructs amenable for automation and parallelization. APL Bioengineering, 4(4). 10.1063/5.0019766

23 Schneider, S., Bubeck, M., Rogal, J., Weener, H. J., Rojas, C., Weiss, M., Heymann, M., van der Meer, A. D., & Loskill, P. (2021). Peristaltic on-chip pump for tunable media circulation and whole blood perfusion in PDMS-free organ-on-chip and organ-disc systems. Lab on a Chip, 21(20), 3963–3978. 10.1039/d1lc00494h

24 Frey, O., Misun, P. M., Fluri, D. A., Hengstler, J. G., & Hierlemann, A. (2014). Reconfigurable microfluidic hanging drop network for multi-tissue interaction and analysis. Nature Communications, 5(1). 10.1038/ncomms5250

25 ISO 22916:2022(en), Microfluidic devices — Interoperability requirements for dimensions, connections and initial device classification

26 Picollet-D’hahan, N., Zuchowska, A., Lemeunier, I., & Le Gac, S. (2021). Multiorgan-on-a-chip: A systemic approach to model and Decipher Inter-organ communication. Trends in Biotechnology, 39(8), 788–810. 10.1016/j.tibtech.2020.11.014

27 SBS Recommended Microplate Specifications

28 Dekker, S., Buesink, W., Blom, M., Alessio, M., Verplanck, N., Hihoud, M., Dehan, C., César, W., Le Nel, A., van den Berg, A., & Odijk, M. (2018). Standardized and modular microfluidic platform for Fast Lab on Chip System Development. Sensors and Actuators B: Chemical, 272, 468–478. 10.1016/j.snb.2018.04.005

29 Vivas, A., van den Berg, A., Passier, R., Odijk, M., & van der Meer, A. D. (2022). Fluidic circuit board with modular sensor and valves enables stand-alone, tubeless microfluidic flow control in organs-on-chips. Lab on a Chip, 22(6), 1231–1243. 10.1039/d1lc00999k

30 de Graaf, M. N., Vivas, A., Kasi, D. G., van den Hil, F. E., van den Berg, A., van der Meer, A. D., Mummery, C. L., & Orlova, V. V. (2023). Multiplexed fluidic circuit board for controlled perfusion of 3D blood vessels-on-a-chip. Lab on a Chip, 23(1), 168–181. 10.1039/d2lc00686c

31 Vollertsen, A. R., de Boer, D., Dekker, S., Wesselink, B. A., Haverkate, R., Rho, H. S., Boom, R. J., Skolimowski, M., Blom, M., Passier, R., van den Berg, A., van der Meer, A. D., & Odijk, M. (2020). Modular operation of microfluidic chips for highly parallelized cell culture and liquid dosing via a fluidic circuit board. Microsystems & Nanoengineering, 6(1). 10.1038/s41378-020-00216-z

32 E. Safai, “Translational Organ-on-chip Platform (TOP) Design Rules (TDRs)”, 2024. doi: 10.4121/2558bd4c-d7ad-4e17-bc54-8c335b4c1c01

33 Eslami Amirabadi, H., Donkers, J. M., Wierenga, E., Ingenhut, B., Pieters, L., Stevens, L., Donkers, T., Westerhout, J., Masereeuw, R., Bobeldijk-Pastorova, I., Nooijen, I., & van de Steeg, E. (2022). Intestinal explant barrier chip: Long-term intestinal absorption screening in a novel microphysiological system using tissue explants. Lab on a Chip, 22(2), 326–342. 10.1039/d1lc00669j

34 Emmerich, M., Ebner, P., & Wille, R. (2024). Automated Design for Multi-Organ-on-chip geometries. IEEE Transactions on Computer-Aided Design of Integrated Circuits and Systems, 1–1. 10.1109/tcad.2024.3509795

35 Biology-inspired dynamic microphysiological system approaches to revolutionize basic research, healthcare and Animal Welfare. (2025). ALTEX. 10.14573/altex.2410112

36 Mastrangeli, M., & van den Eijnden-van Raaij, J. (2021). Organs-on-chip: The way forward. Stem Cell Reports, 16(9), 2037–2043. 10.1016/j.stemcr.2021.06.015

37 Donkers, J. M., Wiese, M., van den Broek, T. J., Wierenga, E., Agamennone, V., Schuren, F., & van de Steeg, E. (2024). A host-microbial metabolite interaction gut-on-a-chip model of the adult human intestine demonstrates beneficial effects upon inulin treatment of gut microbiome. Microbiome Research Reports, 3(2). 10.20517/mrr.2023.79

