## Supplemental Document for "STARTER : A Stand-Alone Reconfigurable and Translational OoC Platform based on Modularity and Open Design Principles"

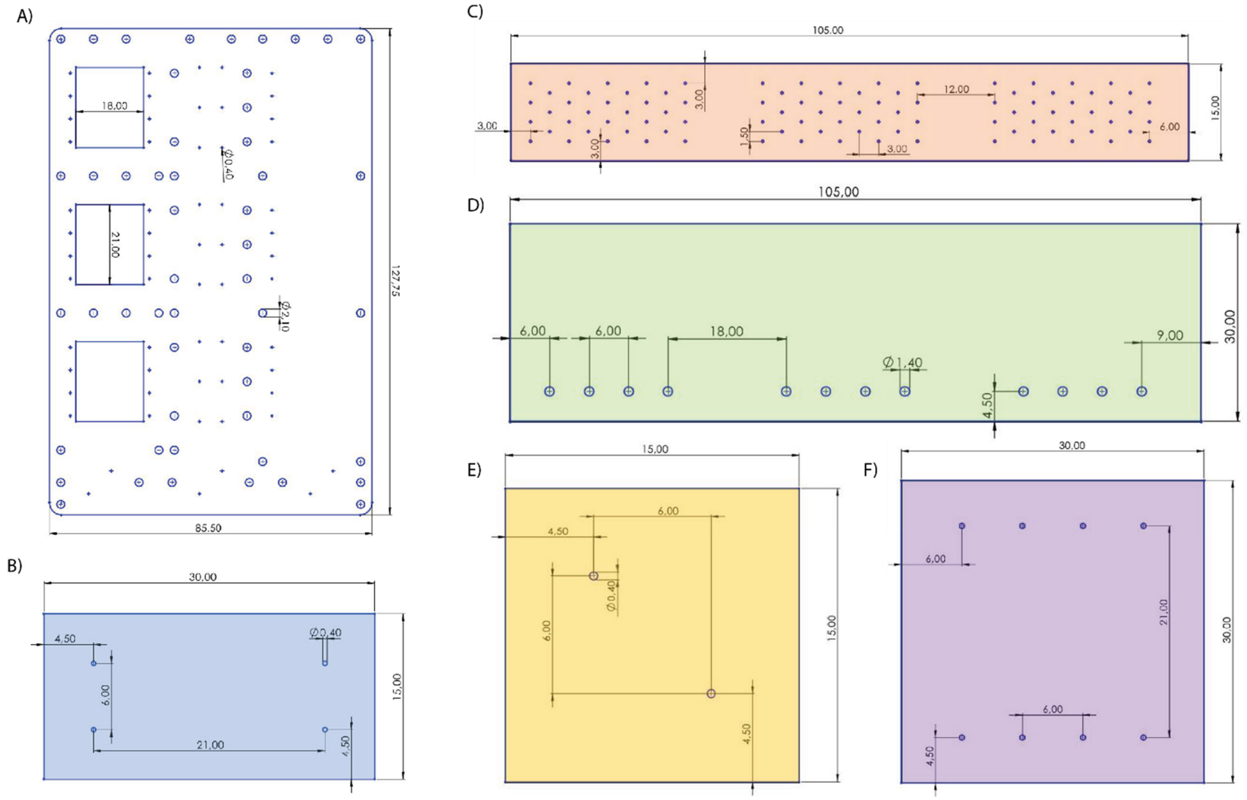


Figure S1 – Ports and Dimensions of STARTER components. a) FCB. b) Reservoir Block. c) Routing Block. d) Pump Block. e) Sensor Block. f) Organ-on-Chip

All MFBBs are designed according to ‘TOP Design Rules’ .

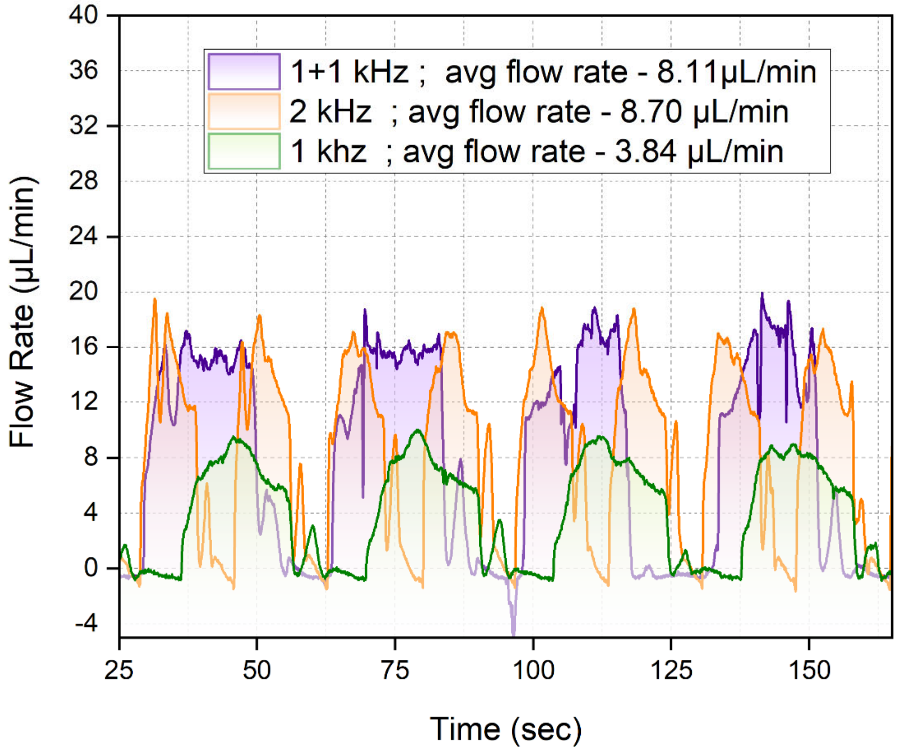


Figure S2 – Flow rate over time comparison between isolated pump operations at 1kHz and 2kHz and combined operation of two pumps running at 1kHz each.

Workflow of STARTER

**Step 1 – OoC cell seeding**. OoCs are seeded off platform till maturation.

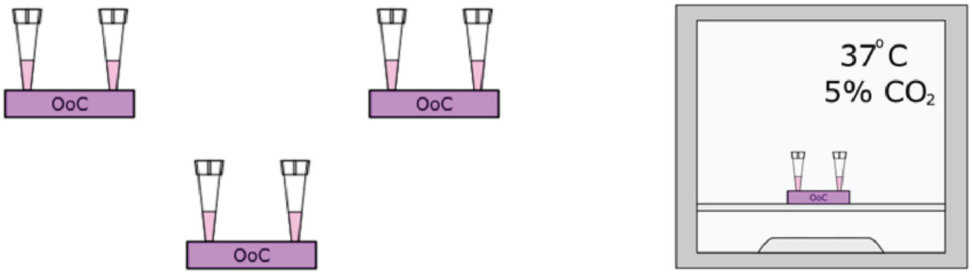


**Step 2 – Sterilizing and priming STARTER.** Dummy priming chips are connected in place of OoCs. The platform is sterilized by flushing with 70% ethanol for 30 min followed by flushing phosphate buffered saline(PBS) for 30 min in the incubator.

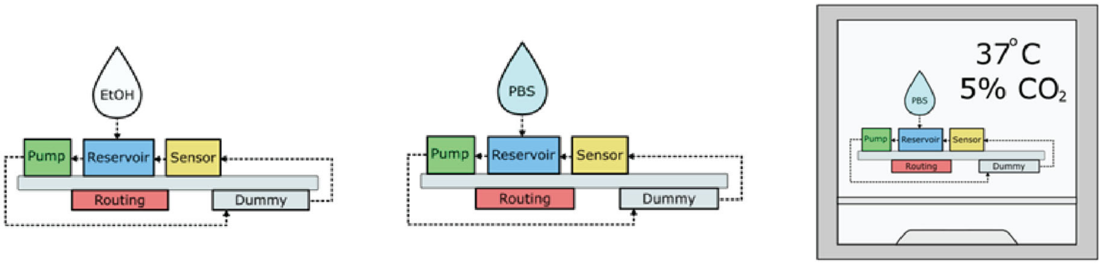


**Step 3 – OoC integration on STARTER**. STARTER is flushed with acclimatized media. The dummy priming chips are then swapped with previously seeded OoCs. The OoCs on STARTER are recirculated with media for required days in incubator.

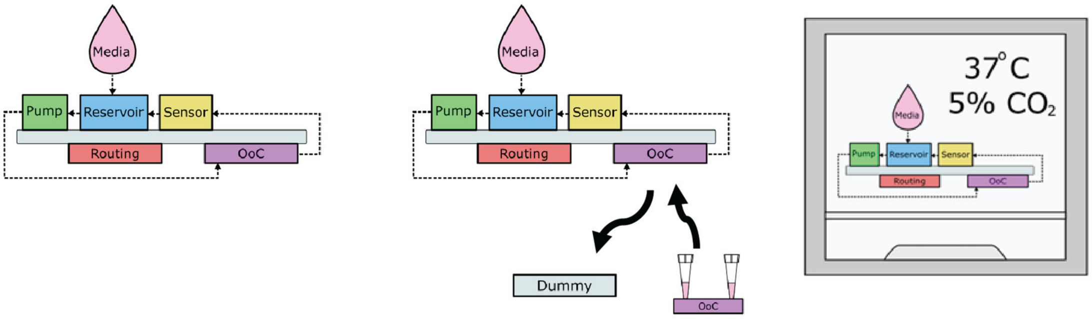

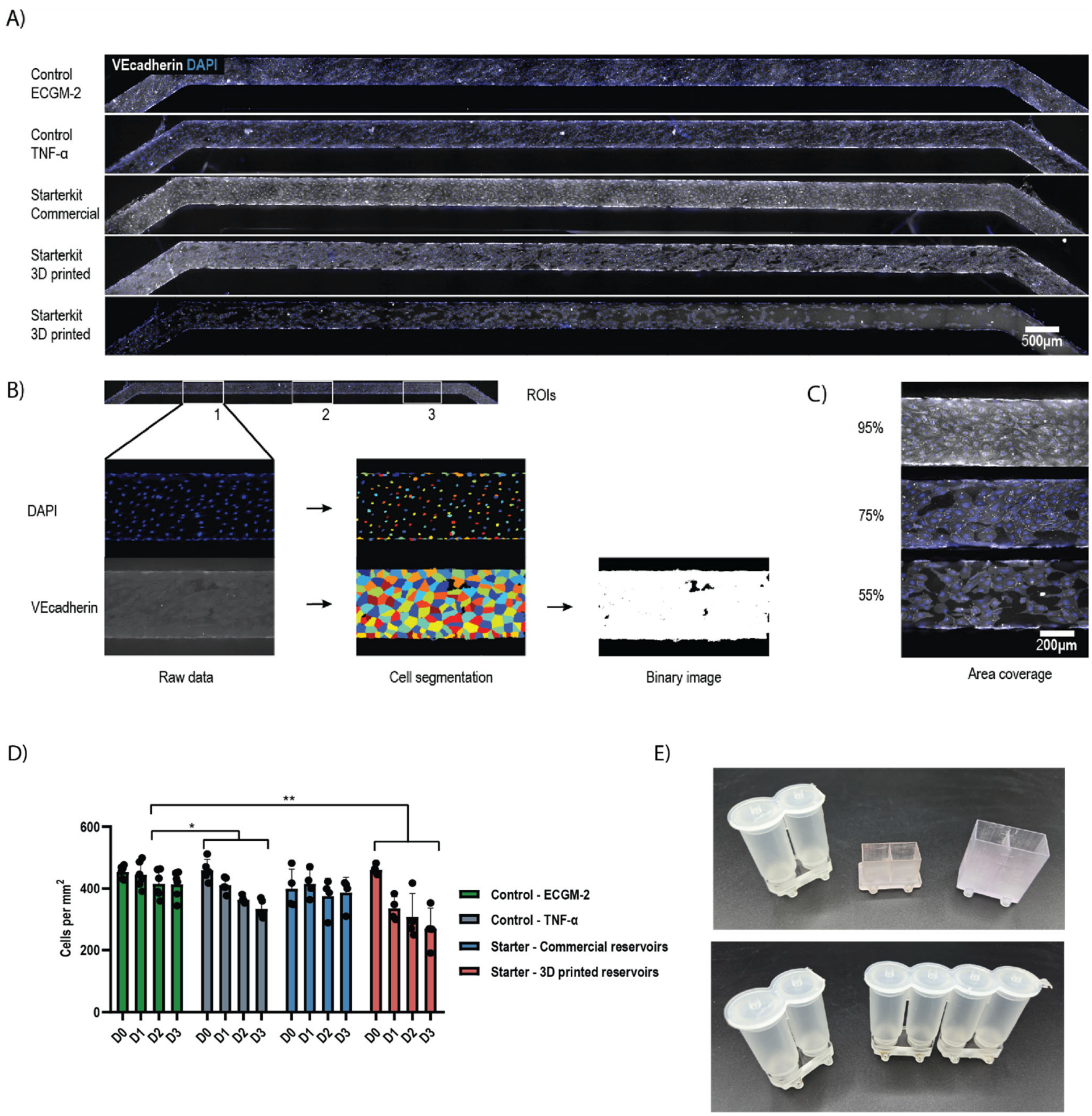


Figure S4 – a) Area coverage of HUVECs after 3 days of cell culture in different conditions. b) Analysis of area covered by cells. c) Example of 95%, 75% and 55% area coverage. d) Cell number during 3 days of in vitro cell culture of HUVECs in a vessel-on-chip (VoC) in regular medium (ECGM-2). Decline in cell numbers in STARTER with 3D printed reservoirs is comparable with pro-inflammatory stimulus with 5ng/ml TNF-α. e) Images of Commercial and 3D printed reservoirs.

*Medium refreshment was either performed on a rocker platform or continuously for control and STARTER conditions, respectively.

### Supplementary Videos-

S1 - Filling of STARTER in Combined Operation

S2 – Mixing and gradient generation
